# Oligomerization enhances huntingtin membrane activity but is suppressed by covalent crosslinking

**DOI:** 10.1101/2023.03.01.530665

**Authors:** Faezeh Sedighi, Adam Skeens, Adewale Adegbuyiro, John Bard, Chathuranga Siriwardhana, Emily Donley, Werner J. Geldenhuys, Justin Legleiter

## Abstract

Huntingtin disease (HD) is a neurodegenerative disease caused by expansion of a polyglutamine (polyQ) tract within the huntingtin (htt) protein, leading to aggregation into a variety of species ranging from small oligomers to large fibrils. A consensus concerning which of these aggregate states are primarily responsible for toxicity associated with mutant htt remains elusive. Htt directly binds and damages a variety of membranous surfaces within cells. Here, the ability of different aggregation states of htt to interact with and damage lipid membranes was determined. Oligomers represented the most active lipid binding species, whereas, fibril formation severely limited membrane binding. Thus, strategies to stabilize oligomers were implemented, and conformational flexibility appeared to play a key role in the oligomer/membrane interaction. In particular, stabilizing oligomers with covalent crosslinking with 1,5-difluoro-2,4-dinitrobenzene (DFDNB) effectively eliminated the ability of oligomers to bind lipid membranes and reduced their associated cellular toxicity.

## INTRODUCTION

Huntington’s disease (HD), a fatal neurodegenerative disorder, is one of many disorders in which an expanded polyglutamine (polyQ) tract leads directly to protein aggregation.^1, 2^ The polyQ expansion (beyond ∼35 repeat units) associated with HD occurs within the first exon of the huntingtin protein (htt).^3^ Mutant htt deposits as intranuclear and cytoplasmic inclusion bodies composed of fibrillar htt aggregates.^4, 5^ Beyond inclusions, htt forms numerous oligomeric and fibrillar species diffusely distributed throughout cells,^2, 6-11^ resulting in a complex mixture of distinct aggregate species and making it difficult to gain consensus on the prominent aggregate species involved in toxicity. Inclusion formation correlates poorly with toxicity,^12^ and survival analysis of cultured neurons suggested inclusion formation is protective, as maintaining a cellular diffuse population of htt correlates strongly with risk of death.^13^ The diffuse cellular population of htt contains a heterogeneous mixture of monomers, oligomers,^8, 14, 15^ and fibrils.^10, 16^ Diffuse aggregates can range in size from small dimers and tetramers^9^ up to ∼1000–3000 molecules.^7^ Complicating matters, toxicity is attributable to monomeric polyQ peptides,^17^ htt oligomers,^15, 18, 19^ and fibrils.^20, 21^ As such, different aggregates may contribute to unique toxic mechanisms within HD.

Htt localizes to a variety of membranous surfaces and organelles, including mitochondria, the endoplasmic reticulum (ER), the nuclear envelope, tubulovesicles, endosomes, lysosomes, presynaptic and clathrin-coated vesicles, and the dendritic plasma membrane.^5, 22-28^ These htt/lipid interactions often result in membrane abnormalities and disruption.^22, 23, 29, 30^ In both HD patients and mouse models, mutant htt exasperates age-dependent nuclear envelope disruption, invoking DNA damage.^23, 31^ Furthermore, htt fibrils impinge on cellular endomembranes, damaging their integrity and freezing ER dynamics.^22^ In mammalian cell and primary neuron models, cytoplasmic htt inclusions contain remnants of organelle membranes from the ER and mitochondria, which correlates with impaired organelle function and localization.^32^

In htt-exon1, the polyQ domain is preceded on its N-terminal side by a seventeen amino acid sequence (Nt17) that is heavily involved in oligomer formation and fibril nucleation.^33-35^ Nt17 appears intrinsically disordered.^33, 36^ but can adopt an amphipathic α-helical structure.^33, 37, 38^ This ability to form an amphipathic α-helices allows Nt17 to function as a lipid binding domain.^36, 39-42^ Htt/lipid interactions further modify aggregation, promoting distinct aggregation pathways in a membrane composition dependent manner.^43-46^

While these observations support a role for htt/lipid interactions in modifying the aggregation process and HD pathogenesis, little is known concerning how aggregation state impacts the membrane activity of htt. Here, the membrane activity of distinct htt aggregates was assessed. These analyses support the notion that oligomerization enhances the direct interaction between htt and lipid membranes; however, fibrillization suppresses membrane activity. Within Nt17, there are three lysine residues that influence oligomer formation and lipid binding.^47-49^ As a result, these lysine residues were targeted by 1,5-difluoro-2,4-dinitrobenzene (DFDNB) to stabilize htt oligomers. While sufficiently large doses of DFDNB stabilized htt oligomers, this negated their ability to interact with lipid vesicles and reduced their toxicity to cell culture, suggesting that reduced flexibility within oligomers underscores their ability to damage membranes.

## METHODS

### Expression and purification of GST-htt-exon1 fusion protein

Glutathione S-transferase (GST)-htt exon1 fusion proteins with a pathogenic (46Q) or nonpathogenic (20Q) length polyQ domain were expressed and purified as previously described. ^50^ Briefly, the protein was expressed in *Escherichia coli* for 4 h at 30 °C and lysed. Proteins were purified using a BioRad low pressure liquid chromatograph equipped with a GST-affinity column. The purity and the relevant fractions were verified by SDS-PAGE. Relevant fractions were pooled and placed in dialysis at 4 °C for 2 days. To remove potential pre-aggregated species, the protein solution was centrifuged at (22,000 ×g for 30 min) before any experiments. Protein concentration was determined by a Bradford assay. To initiate aggregation, the desired volume of protein solution was incubated with Factor Xa (Promega, Madison, WI). All experiments were carried out in a tris buffer (150 mM NaCl, Tris-HCl, pH 7.4).

### Atomic force microscopy (AFM)

*Ex situ* atomic force microscopy (AFM) was used to characterize the morphology of htt aggregates. For pre-aggregation experiments, htt-exon1(46Q) or htt-exon1(20Q) (10 µM) were incubated 30 °C for and sampled after 1, 3, 5, and 8 h by taking a 3 µL aliquot and depositing it on freshly cleaved mica for 1 min. Then, the mica was rinsed with 150 µL of ultrapure water and dried with a gentle stream of air. For morphological analysis of crosslinked htt, aliquots were taken before crosslinking, immediately after crosslinking, and 24 h after crosslinking. For morphological analysis of oligomers formed in the presence of Nt17 peptides, samples were taken after 3 h of incubation and after an additional 24 h. Samples were imaged using a Nanoscope V Multi-Mode scanning probe microscope (VEECO) equipped with a closed-loop vertical engage J-scanner. Silicon-oxide cantilevers with nominal spring constant of 40 N/m and a resonance frequency of 300 kHz were used. Scan rates were set to 1.99 Hz with cantilever drive frequencies at 10% off resonance.

For *in situ* AFM experiments that tracked the interaction of htt with supported total brain lipid extract bilayers (TBLE), a tapping mode fluid cell equipped with an O ring and a cantilever with nominal spring constant of 0.1 N/m was used. The TBLE bilayer was prepared by reconstituting a lyophilized lipid film in the Tris buffer to a concentration of 1 mg/mL. The lipid solution then underwent 10 freeze-thaw cycles using liquid nitrogen followed by bath sonication for 10 minutes. The resulting lipid vesicle solution was injected into the fluid cell (30 µL), and the formation of the lipid bilayer was monitored by continuous AFM imaging (∼2 h) until a complete 40 µm × 40 µm supported bilayer was formed. The bilayers were exposed to preformed oligomers, fibrils, or crosslinked oligomers by direct injection (20 µM of 30 µL, resulting in a final concentration of 10 µM htt) into the fluid cell. Oligomers and fibrils were prepared by 3 and 24 h of pre-incubation and verified via *ex situ* AFM. Crosslinked oligomers were prepared as described below. Morphological changes on the lipid bilayer were monitored by continuous imaging for 3 h.

To further assess the aggregate populations and morphological changes in AFM images (both *ex situ* and *in situ*), automated algorithm written in Matlab measured morphological features of all features in AFM images,^51^ Oligomers were defined as particle taller than 1 nm with an aspect ratio of less than 2.5, indicating a globular structure. Due to extensive bundling, the fibril population was determined by visual inspection of all images.

### Polydiacetylene lipid binding assay

Normalized polydiacetylene (PDA) lipid binding assays were utilized to measure the interaction between htt-exon1 (both 46Q and 20Q) and TBLE. Briefly, diacetylene monomers (10,12-tricosadiynoic acid) and TBLE were dissolved in a 4:1 chloroform:ethanol solution and mixed at a 2:3 molar ratio. Solutions were dried under a gentle stream of nitrogen until a fully dried lipid film was formed. These films were reconstituted in hot tris buffer (150 mM NaCl, 50 mM Tris-HCl, pH 7.4, 70 °C), sonicated at 125 W for 10 min, protected from light, and left at 4 °C overnight to allow vesicle formation. The following day, the diacetylene monomers in the lipid vesicles were polymerized by exposure to UV light (254 nm), resulting in a royal blue solution that undergo a colorimetric shift to red upon mechanical stress. PDA/TBLE solutions (400 µM) were incubated with different preparations of htt-exon1(46Q) at 30 °C, and the blue (650 nm) and red (500 nm) absorbance (A_blue_ and A_red_ respectively) were recorded every 5 min for 12 to 20 h. The negative control consisted of equal volumes of neat buffer and PDA/TBLE solution, while the internal standard involved equal volumes of 1M NaOH and PDA/TBLE. The NaOH creates repulsion between headgroups via protonation, causing a saturated colorimetric shift that can be used to normalize results.^52^ Additional controls with corresponding DFDNB/DTT concentration for each condition were also performed. The % CR was calculated for each condition using the following equation:

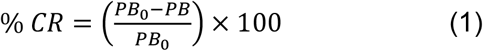

where PB is defined as A_blue_/(A_blue_ + A_red_) for the negative control (PB_0_) and sample condition (PB). ^53, 54^

### Crosslinking htt with DFDNB

For crosslinking oligomers, htt-exon1(46Q) and htt-exon1(20Q) was incubated (after factor Xa treatment) at 40 µm for 1 and 3 h on ice. Next, the htt samples were treated with varying concentration (2.5 to 20 fold excess relative to final htt concentration of 20 µM) of DFDNB. After mixing by inversion, the crosslinking interaction ran for 10 min at 20 ºC. After which, the reaction was quenched by equimolar concentration of dithiothreitol (DTT), and allowed to sit for 10 min The resulting solutions were used in PDA, ThT, or MTT assays. All crosslinked htt reactions were characterized by *ex situ* AFM analysis.

### Thioflavin-T aggregation assay (ThT)

Thioflavin T fluorescence assays (ThT, Sigma-Aldrich, St. Louis, MO) were performed to monitor fibril formation. Experiments were performed in multi-well plate format (Costar clear-bottom 96-well plates) using a SpetraMax M2 microplate reader (440 nm excitation, 484 nm emission) with readings taken every 5 min for 15 h. The ThT concentration in each well was 200 µM. Experiments were performed in Costar clear-bottom 96-well plates at 37 °C and ThT. Each experiment was performed three separate times, and within each experiment each condition was run in triplicate. Controls included neat buffer and corresponding concentration of DFDNB/DTT for each condition used for oligomer crosslinking.

### Dynamic light scattering

Dynamic light scattering (DLS) was used to measure the size distribution of the htt control and crosslinked htt after 24 h, 3 days, and 10 days after the crosslinking reaction. DLS experiments were performed on a 90Plus/BI-MAS nanoparticle size analyzer (Brookhaven instrument). The wavelength of the laser was 635 nm and the scattered light was measured at 90°. Htt samples were diluted to a final concentration of 4 µM. Due to the formation of larger aggregates in the htt control (non-crosslinked) samples and crosslinked htt with 5 x DFDNB treatment, a high speed centrifuge step (5,000 g) for 20 min was required to pellet the larger size particles and allow the suspensions to be in the size limit range of the instrument. Measurements were taken at 25 °C, and the data was analyzed using the MAS OPTION software.

### Mtt toxicity assay

The cytotoxicity of htt oligomers, DFDNB crosslinker and, crosslinked oligomers was tested on the Neuro-2a (N2a) cell line (ATCC® CCL-131™) using Mitochondrial 3-(4,5-dimethylthiazol-2-yl)-2,5-diphenyltetrazolium bromide (MTT) reduction assay. Oligomers were prepared by 3 h incubation of htt-exon1(46Q) on ice at a concentration of 20 μM. Some oligomers were crosslinked with 20× or 5× DFDNB treatments under the same condition described before. The crosslinked reactions were quenched by the equimolar concentration of DTT after 10 min. The final concentration of the oligomers in the crosslinked solutions were 10 μM. Vehicle controls of DFDNB corresponding to 200 μM were prepared to measure the toxic potency of the free DFDNB crosslinker and all chemical treatments required for crosslinking. To reduce the excess free DFDNB molecules, crosslinked oligomers and DFDNB vehicle controls were dialyzed for 24 h with (300 μL of sample to 10 mL of dialysis buffer). On the day of experiment, fresh htt oligomers at a concentration of 10 μM were used as a control. Stabilized oligomers, fresh non-crosslinked oligomer controls, and DFDNB vehicle controls were added to the extracellular medium of cultured N2a cells plated in triplicate in a flat-bottom 96-wells microplate. The samples were added via serial dilution resulting in doses ranging from 0.2 μM to 5 μM final concentration of htt in the wells. The DFNDB concentation of the vehicle control in the wells varied from 0.2-0.08 μM based on equivalent volume added for the htt samples. To facilitate transferring the oligomers, cells were permeabilized with 0.1% saponin prior to the experiment, and the MTT assay was performed 72 h after the addition of htt. At this time, the MTT assay was performed as described in the MTT assay kit protocol (Millipore Sigma).

## RESULTS

### Htt oligomer formation correlates with lipid binding

The ability of htt-exon1 to directly bind lipid membranes is well established, yet little is known concerning the role of aggregation state on this process. To gain an initial understanding of how aggregation state impacts htt/lipids interactions, 10 µM htt-exon1(46Q) samples were pre-incubated at 30 °C for variable lengths of time (1, 3, 5, and 8 h), producing different populations of htt aggregates that were analyzed via *ex situ* AFM (Fig 1A-B). The number of oligomers and fibrils per µm^2^ was determined (Figure 1B). A substantial number of oligomers were observed after 1 h of incubation (17.0 ± 1.5 per µm^2^), and the oligomer population peaked (23.2 ± 1.4 per µm^2^) after 3 h. With longer incubation times, the oligomer population steadily decreased (17.9 ± 2.5 per µm^2^ and 8.4 ± 3.1 per µm^2^ at 5 and 8 h respectively). This decrease in oligomer population occurred concurrently with the appearance (at 5 h) and subsequent (at 8 h) of fibrils. Morphological differences were observed in oligomers with time. After 1, 3, and 5 h of incubation, oligomers had a mode height of ∼3.5-4.5 nm and a mode diameter of ∼60-70 nm (Figure 1C). However, a distinct tail in the diameter histogram developed with time. After 8 h, oligomers were clearly larger (mode height of ∼5-6 nm and a mode diameter of ∼80-100 nm), signifying that oligomer morphology changed with time.

**Figure 1.**
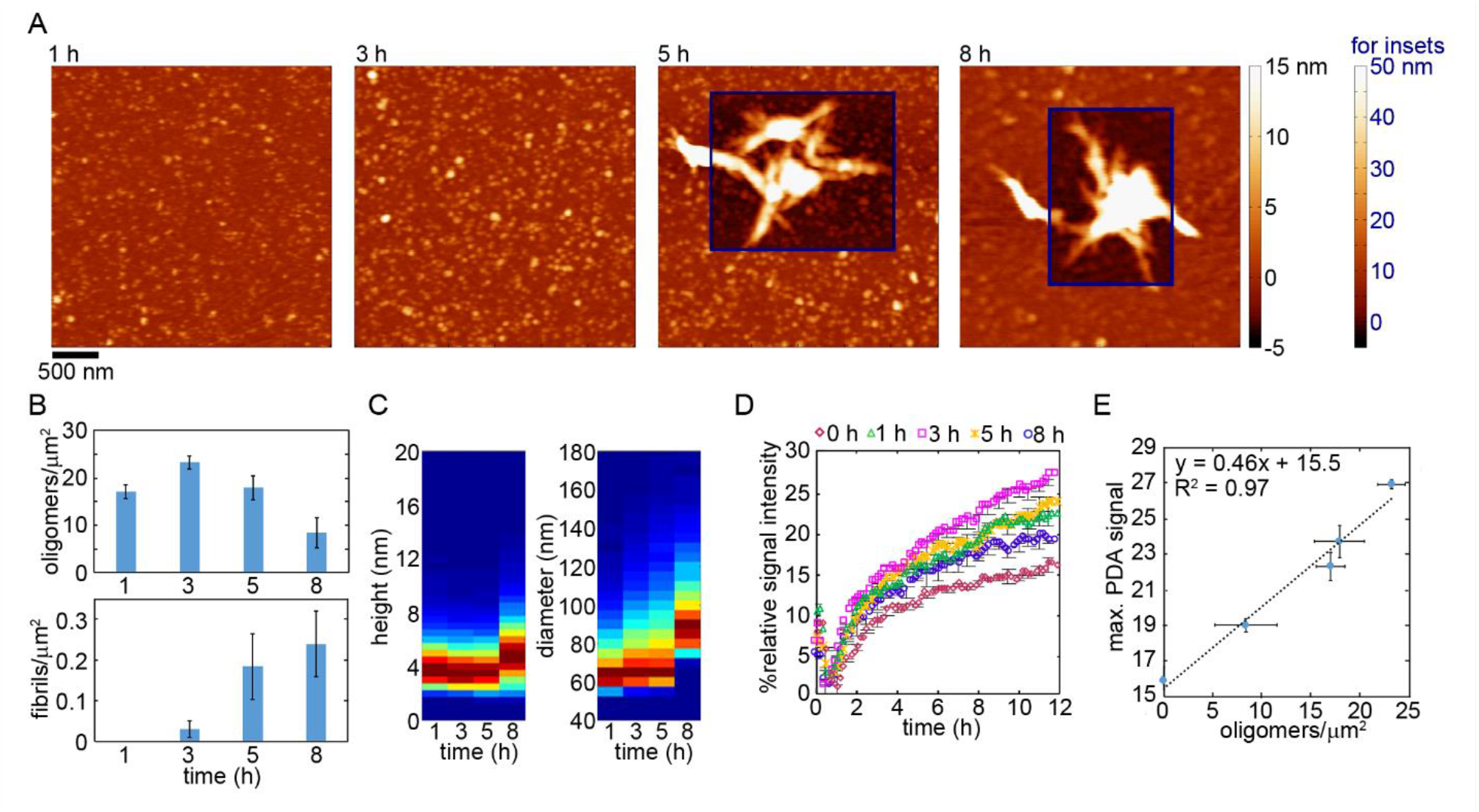
(A) Representative *ex situ* AFM images of htt-exon1(46Q) aggregates present after 1, 3, 5, or 8 h of incubation at 10 µM. The second color bar (labeled in blue) corresponds to the insets in the images marked by blue boxes. (B) Analysis of oligomer and fibril populations present at each time point. (C) Histograms of oligomer height and diameter as a function of time. (D) PDA/TBLE lipid binding assay performed with htt-exon1(46Q) pre-incubated for various length of time. (E) The maximum PDA signal plotted as a function of the oligomers per unit area observed by AFM. Error bars represent standard deviation taken across multiple images and experiments.

The extent to which these different aggregate populations bound a total brain lipid extract (TBLE) bilayer was determined via a polydiacetylene (PDA) lipid binding assay (final htt concentration of 5 µM, Figure 1E). Mechanical stress applied to the PDA backbone of PDA/TBLE vesicles by protein binding induces a quantifiable colorimetric response (CR). Upon exposure of PDA/TBLE vesicles to freshly cleaved htt-exon1(46Q) (0 h incubation, predominately monomeric), a steady increase in the PDA signal was observed, maxing out at ∼ 15.9% relative intensity after 12 h (Figure 1D). The 1 h incubation that contained oligomers elicited a larger CR, reaching 22.3% relative intensity. The 3 h incubation, containing the largest oligomer population, invoked the largest PDA signal with a maximum value 26.9% relative intensity. The PDA signal produced by the 5 h incubation, which contained fibrils, was less intense compared to the 3 h incubation, maxing out at 23.7%. Lastly, the 8 h htt incubation, which contained fewer oligomers and more fibrils, resulted in an even greater reduction in the PDA signal (maximum of 19.0%). The maximum PDA signal correlated linearly with the oligomer population size (Figure 1E, R^2^ = 0.97), suggesting that oligomerization enhances the ability of htt to bind membranes.

A complication of the previous analysis is that these incubations still contained complex mixtures of htt species. That is, as fibrils were not the predominate aggregate in any of the incubation, the relative fibril affinity for membranes was difficult to assess. Therefore, a PDA/TBLE assay was performed with freshly prepared (predominately monomeric) htt, oligomers, and fibrils at a final concentration of 10 µM for all three conditions (Figure 2). To obtain oligomers, htt was incubated on ice for 3 h. To obtain fibrils, htt-exon1(46Q) was seeded with preformed fibrils (0.25 µM concentration) of a polyQ peptide and incubated for 24 h at 30°C. Prior to the PDA/TBLE assay, aggregate type was verified by AFM (Figure 2A). The freshly prepared samples contained a few oligomers but significantly fewer compared to the oligomer prep. The fibril sample contained long, entangled fibrils. The oligomer prep elicited the largest CR from the PDA/TBLE assay, ∼3-fold increase in maximum signal compared with fresh htt. Fibrils induced the smallest relative CR compared to oligomers (∼45% reduction).

**Figure 2.**
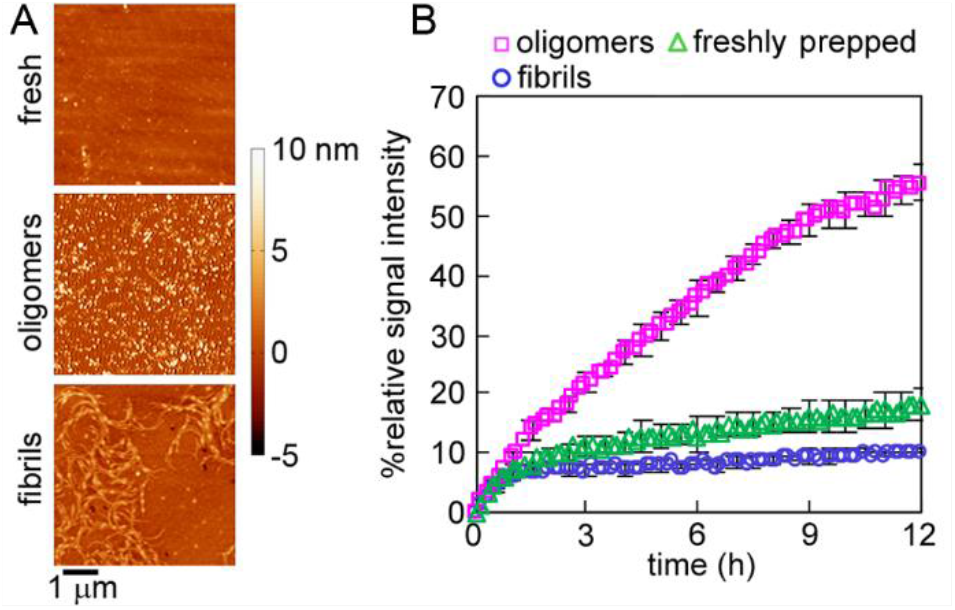
(A) *Ex situ* AFM images of different preparations of htt-exon1(46Q) aggregates (freshly htt, oligomers, and fibrils) that were used for (B) PDA/lipid binding assays.

### Crosslinking alters htt aggregation in a dose-dependent manner

Having established that htt oligomers have a relatively large membrane activity, the possibility of stabilizing oligomers via chemical crosslinking was explored to facilitate further investigating the oligomer/membrane interaction. As Nt17 promotes htt aggregation via an α-helical oligomeric intermediate,^33^ it represents a viable target for oligomer stabilization. There are three lysine residues located within Nt17, two (K6 and K15) of which are implicated in oligomer formation.^47^ As a result, a lysine specific crosslinking agent, 1,5-difluoro-2,4-dinitrobenzene (DFDNB), was employed to crosslink oligomers. DFDNB crosslinking is an established method of stabilizing oligomers of other amyloid-forming proteins.^55^ DFDNB was applied to htt-exon1(46Q) incubations after 1 or 3 h with doses ranging from 2.5× to 20× to the final htt concentration (10 µM). To monitor the ability of the different crosslinking conditions to arrest fibril formation, Thioflavin T (ThT) assays were used starting as soon as the DFDNB crosslinking reaction was quenched (Figure 3A-B). Control htt-exon1(46Q) incubations that were only treated with vehicle (no DFDNB) that had been incubated for 1 h or 3 h invoked a continuous increase in ThT fluorescence over 12-15 h (Figure 3A-B). Crosslinking htt-exon1(46Q) after 1 h of pre-incubation with doses of 2.5× and 5× DFDNB actually accelerated aggregation, but DFDNB treatments of 10×, 15×, or 20× suppressed htt aggregation. When htt was treated with different doses of DFDNB after 3 h of pre-incubation, only the 2.5× treatment enhanced aggregation. Doses of 5× DFDNB or higher suppressed fibrillization; although, a slight increase in ThT signal was observed for 5× and 10× towards the end of the assay.

**Figure 3.**
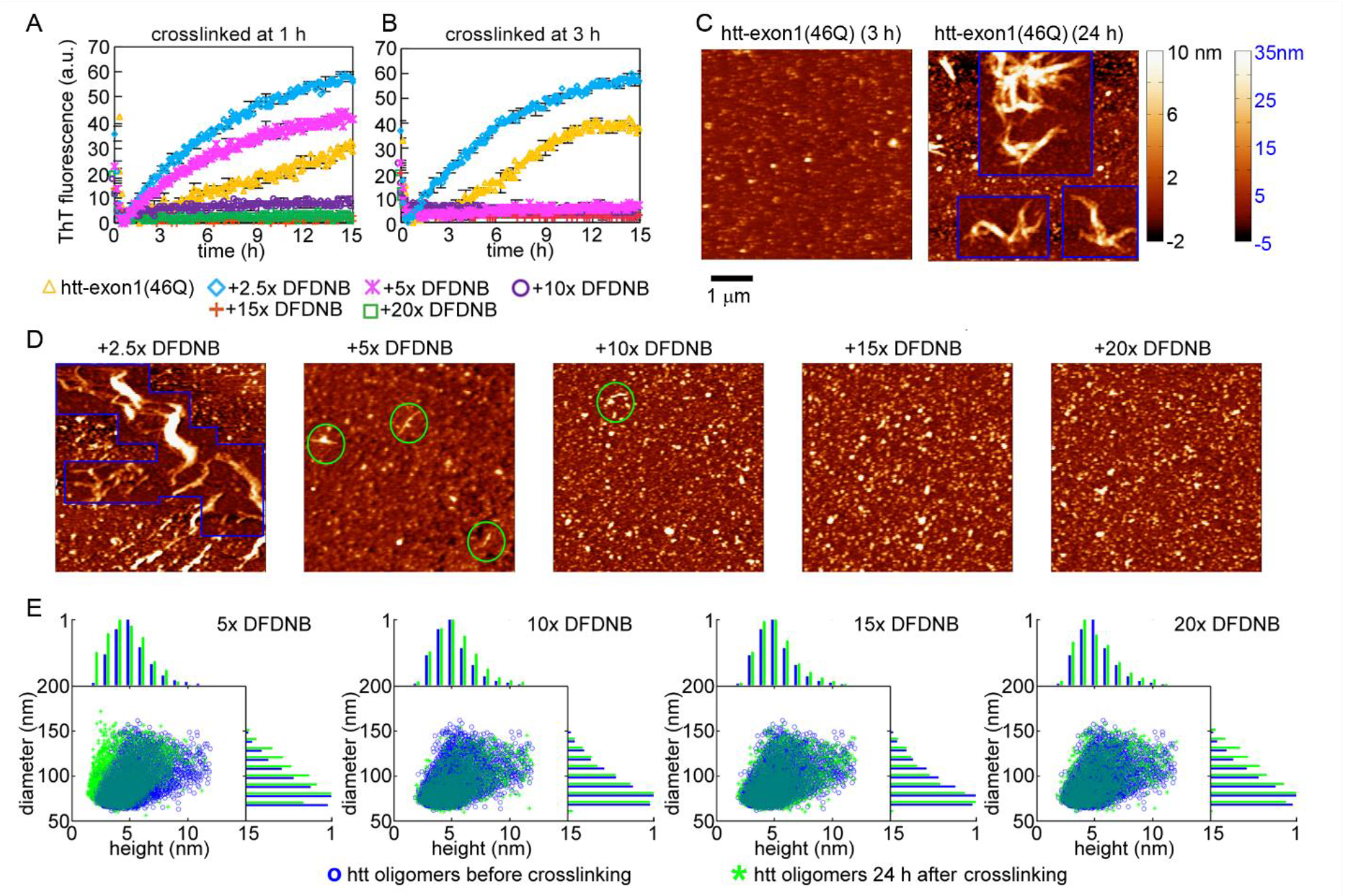
ThT assay tracking aggregation of htt-exon1(46Q) crosslinked after (A) 1 h of pre-incubation or (B) 3 h of pre-incubation. (C) AFM images of htt-exon1(46Q) without crosslinking after 3 and an additional 24 h of incubation. (D) AFM images of htt-exon1(46Q), which was crosslinked with varying doses of DFDNB after 3 h of pre-incubation, imaged after an additional 24 h of incubation. Green circles indicate small protofibrils. For both C and D, the second color bar (labeled in blue) corresponds to the insets in the images marked by blue boxes. (E) Correlation plots of height vs diameter of oligomers taken before crosslinking and an additional 24 h after crosslinking. In addition, the height (top) and diameter (right) histograms are provided.

While ThT assays suggested that large doses of DFDNB suppressed fibrillization, oligomer stabilization was further validated using AFM. Aliquots were taken from untreated and crosslinked htt-exon1(46Q) samples that had been pre-incubated for 3 h immediately after DFDNB treatment and after an additional 24 h of incubation. In the control incubation, abundant oligomers were present at 3 h (Figure 3C). With an additional 24 h of incubation, the control was comprised of oligomers and an extensive network of fibrils. All of the crosslinked samples appeared similar to control at the 3 h time point (large oligomer population). After an additional 24 h, htt-exon1(46Q) treated with 2.5× DFDNB formed a significant fibril population (Figure 3D), consistent with ThT assays. These fibrils were thicker than control fibrils with a slightly curved, less rigid appearance. Larger DFDNB treatments (5× or higher) suppressed fibril formation and appeared to stabilize oligomers; although, a few small protofibrils were observed with 5× and 10× doses of DFDNB after 24 h (Figure 3D). Comparison of oligomer morphology (diameter and height, Figure 3E) indicated that oligomers treated with high doses of DFDNB (10× or higher) were stable. However, oligomers treated with 5× DFDNB had a slight shift to larger heights.

To further validate oligomer stabilization, DLS experiments were performed. Oligomers were crosslinked after 3 h of pre-incubation with 5×, 10×, 15×, or 20× DFDNB, and compared to a control group without any DFDNB added. After 3 days of additional incubation, size distributions of each sample were measured by DLS (Figure 4A). The diameter distributions of control and 5× crosslinked samples were not consistent with oligomers (mode diameter > 1 µm for both). These samples actually required a high-speed spin to pellet larger aggregates so that the remaining aggregates were within the size limits of the instrument. Thus, the reported DLS measurement for this control underestimates the actual size of present aggregates. When crosslinked with DFDNB treatments of 10× or higher, the mode diameter of aggregates was ∼40-70 nm, consistent with the oligomers observed by AFM. These samples treated with 10× DFDNB or higher were incubated for an additional week (10 days after initial crosslinking), and the size distributions for all three samples were stable, validating long term stability of crosslinked oligomers (Figure 4B-D).

**Figure 4.**
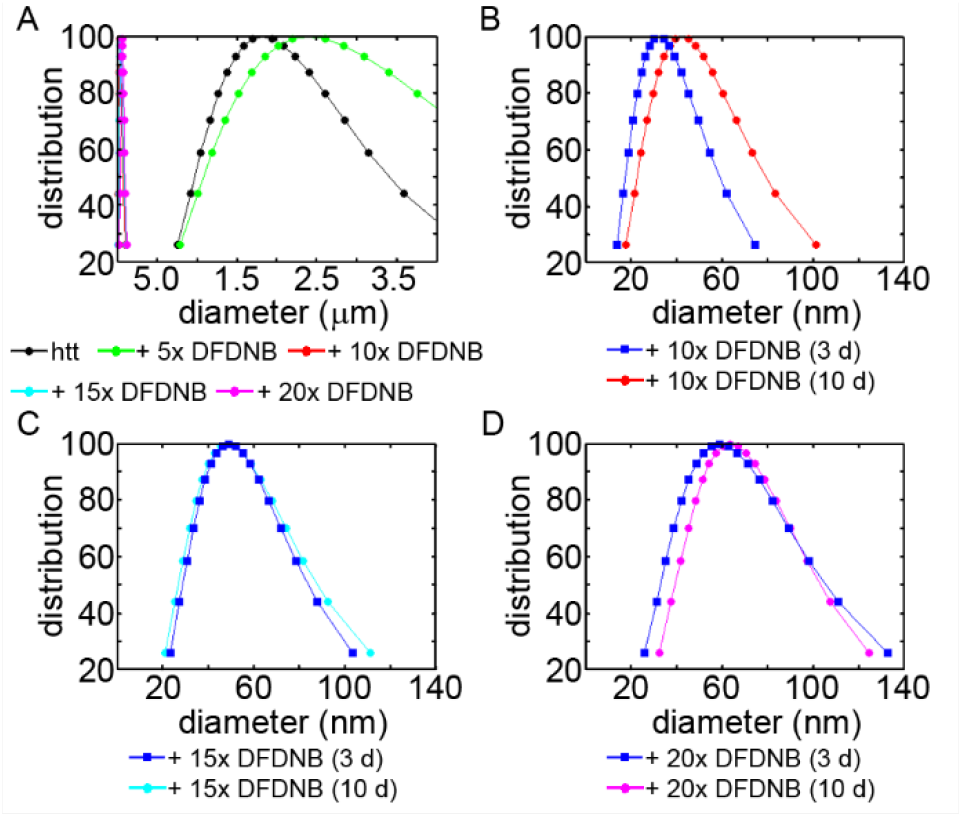
(A) Comparison of aggregate diameter measured by DLS for non-crosslinked and crosslinked htt-exon1(46Q) (after 3 h of pre-incubation) with varying doses of DFDNB after an additional 3 days of incubation. Comparison of aggregate diameter of htt-exon1(46Q) oligomers crosslinked after 3 h of pre-incubation with (B) 10x, (C) 15×, and (D) 20× DFDNB treatments after 3 and 10 days of additional incubation.

### Crosslinking inhibits oligomers from binding lipid vesicles

Having established that htt oligomers had the highest membrane activity in comparison to monomers or fibrils, the membrane activity of DFDNB crosslinked oligomers was determined using PDA/TBLE assays (Figure 5A). After 3 h of incubation, htt-exon1(46Q) oligomers were crosslinked with DFDNB treatments of 2.5× to 20×. As a control, PDA/TBLE vesicles were exposed to untreated htt-exon1(46Q) oligomers, and a large CR was observed (maximum signal of ∼60%). DFDNB treatment reduced the membrane activity of oligomers in a dose-dependent manner. Treatment with 2.5× or 5× DFDNB reduced the maximum CR to ∼54% and 30% respectively. A small CR was associated with oligomers treated with 10× DFDNB, reaching a maximum relative CR of ∼12%. Oligomers stabilized by 15× and 20× had virtually no membrane activity.

**Figure 5.**
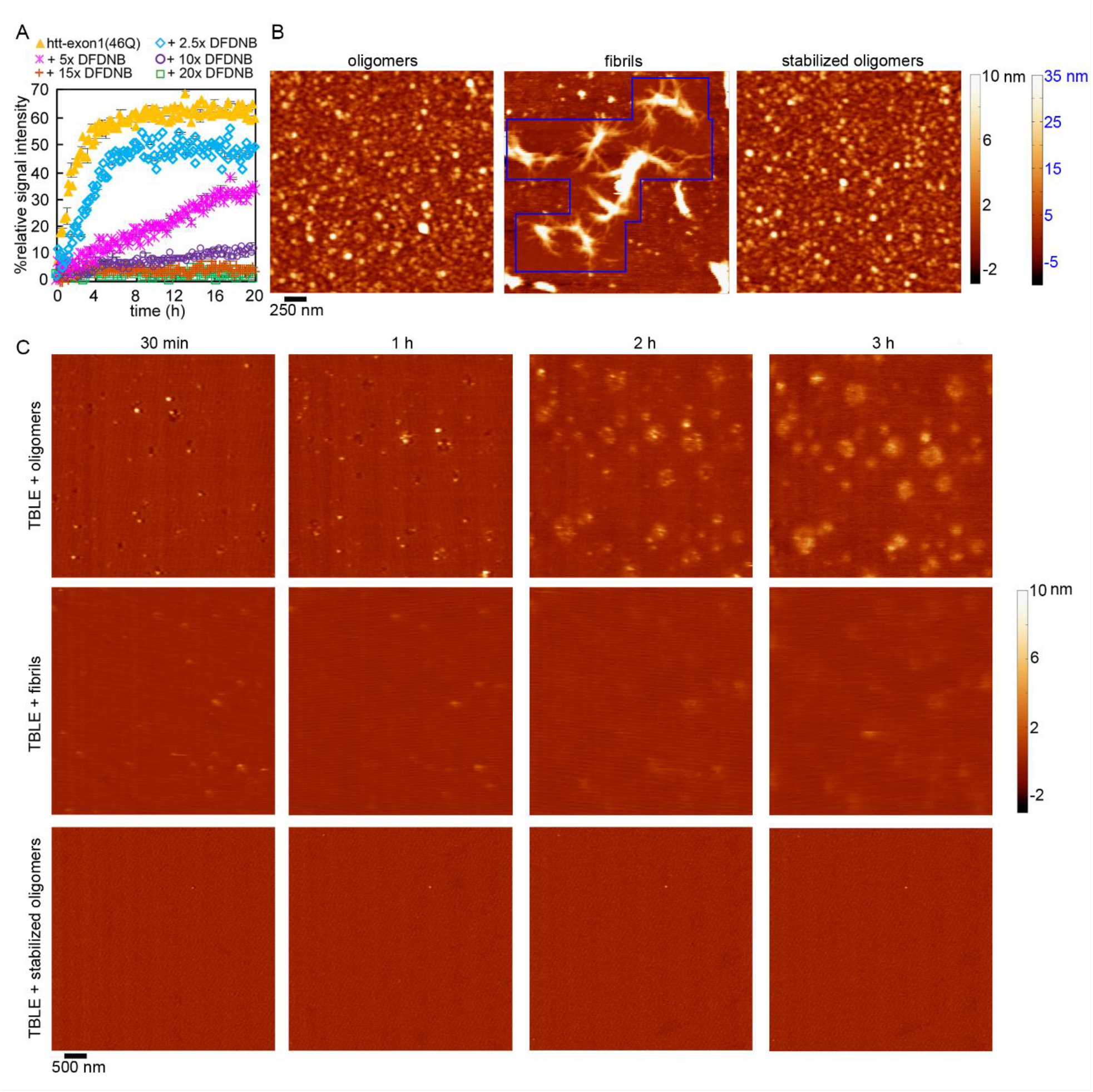
(A) The relative affinity of htt oligomers that were pre-incubated for 3 h without and with varying treatments of DFDNB crosslinking measured by a PDA/TBLE lipid binding assay. *Ex situ* AFM images of htt-exon1(46Q) oligomers (formed after 3 h of incubation), fibrils (formed by seeding and 24 h of incubation), and crosslinked oligomers (3 h of pre-incubation). Time course *in situ* AFM images of a supported TBLE bilayer exposed to htt-exon1(46Q) oligomers, fibrils, and crosslinked oligomers.

To further evaluate the membrane activity of oligomers (both stabilized and untreated) and fibrils, a series of *in situ* AFM was implemented to directly visualize htt aggregates on a lipid bilayer (Figure 5C). Supported TBLE bilayers were exposed to 1) htt-exon1(46Q) oligomers obtained by 3 h of pre-incubation, 2) htt-exon1(46Q) oligomers crosslinked with 20× DFDNB after 3 h of pre-incubation, and 3) htt-exon1(46Q) fibrils (Figure 5B). Untreated oligomers appeared on the bilayer surface within 30 min and were often associated with the development of small holes in the bilayer. With time, regions of increased bilayer roughness developed around oligomers. In the absence of lipids, fibrils are typically observed in incubations of htt-exon1(46Q) at this concentration and time frame (a total of 6 h). However, no fibrils were observed on the bilayer surface, consistent with reports that TBLE inhibits htt fibrillization.^39, 56^ With pre-formed fibrils, bilayer morphology was subtly altered but to a significantly lesser extent in comparison to experiments with untreated oligomers. Again, no fibrils appeared on the surface, but a few oligomers were observed, which may represent residual oligomers left over in the sample prep. When exposing TBLE bilayers to stabilized htt oligomers very few (if any) crosslinked oligomers appeared on the bilayer surface and no membrane roughening occurred over the entire 3 h timeframe.

The disrupted membrane regions invoked by untreated oligomers or fibrils were morphologically different (Figure 6A). Untreated oligomers invoked larger and rougher regions of disruption compared with fibrils. The TBLE bilayer morphology observed upon exposure to DFDNB stabilized oligomers was comparable to freshly formed TBLE bilayers. To quantify this, the RMS roughness of the TBLE bilayer was measured as a function of time. To avoid artifacts in these measurements associated with surface coverage of these regions, two RMS roughness values were determined, disrupted regions (rough) and undisrupted regions (smooth). If perturbed regions were not identified, RMS values were measured across the entire image (Figure 6B). The RMS roughness of a fresh bilayer was ∼ 0.4 nm. Upon exposure to untreated htt oligomers, disrupted regions of RMS roughness of 1.9 ± 0.3 nm appeared at 30 min. The RMS roughness increased with time (2.8 ± 0.9 nm at 3 h). Disrupted bilayer regions associated with fibrils were less pronounced (RMS roughness from 0.6 to 0.9 nm over 3 h). For exposure to both untreated oligomers and fibrils, the RMS roughness of undisrupted (smooth) portions of the bilayer were similar to fresh bilayer. With exposure to DFDNB oligomers, the RMS roughness measured over the entire image for all time points was indistinguishable from a fresh TBLE bilayer. In addition, the area of the bilayer altered by exposure to htt-exon1(46Q) varied by aggregation state (Figure 6C). After 30 min, untreated oligomers disrupted 0.9 ± 0.2 % of the bilayer surface, and the extent of disruption steadily increased (10.7 ± 1.6% after 3 h). Fibrils disrupted the bilayer to a significantly lesser extent, with only 4.0 ± 0.4% of the bilayer being altered within 3 h. Crosslinked oligomers did not induce extensive membrane disruption over 3 h (less than 1% at all time points), and this percentage represents discreet oligomers appearing on the bilayer rather than enhanced roughness.

**Figure 6.**
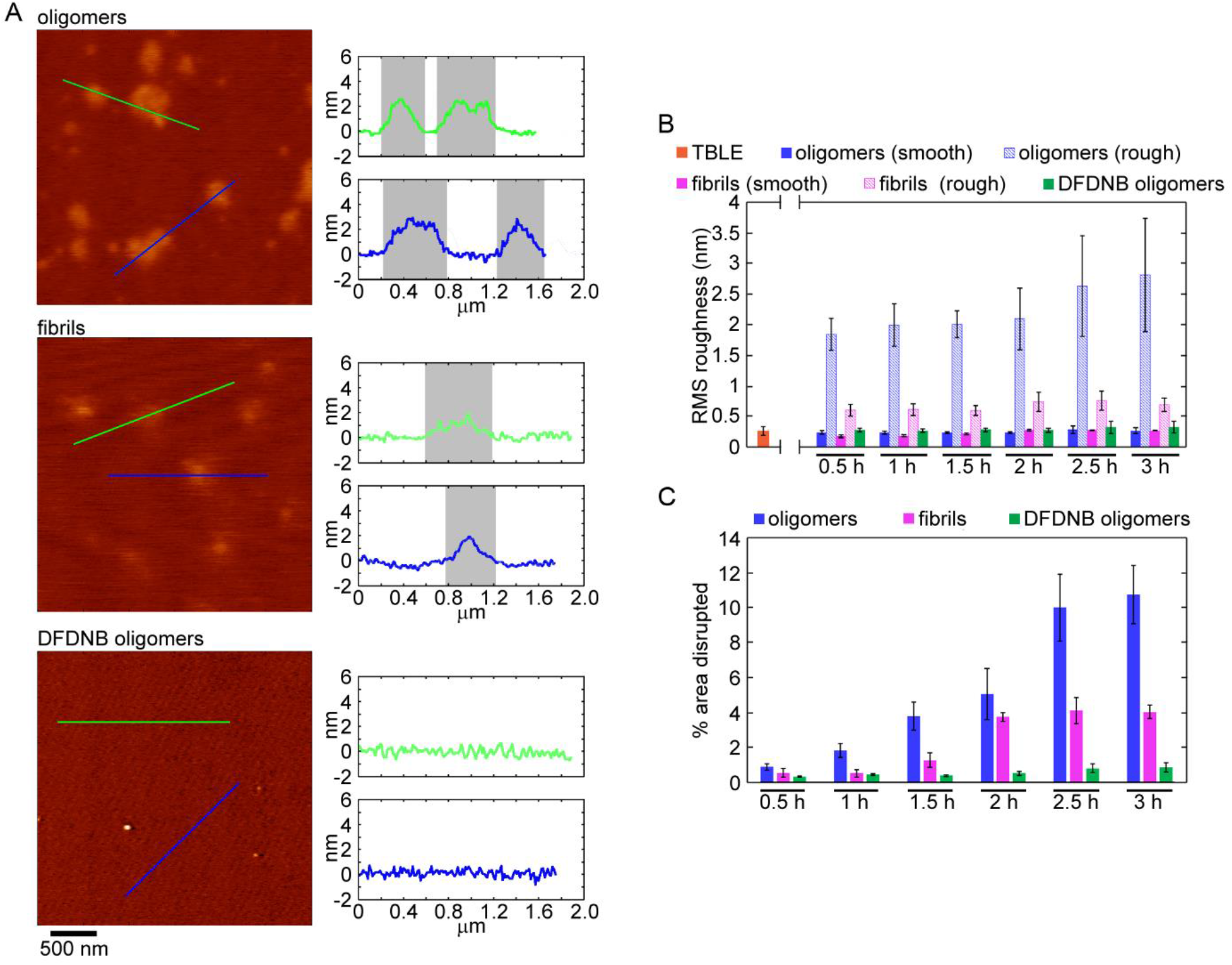
(A) Comparison of the morphology of supported TBLE bilayers exposed to htt-exon1(46Q) oligomers, fibrils, or DFDNB stabilized oligomers. The colored lines in each *in situ* AFM image corresponds to the height profiles provided to the right. In each height profile, the gray shaded areas correspond to regions of the bilayer with increased surface roughness. (B) Comparison of the RMS roughness associated with a TBLE bilayer and the smooth and rough regions observed after exposure to htt oligomers, fibrils, and DFDNB stabilized oligomers as a function of time. As rough regions in the bilayer did not develop upon exposure to DFDNB stabilized oligomers, RMS roughness was measured over the entire image. (C) Comparison of the % area of the TBLE bilayer that was disrupted by exposure to htt oligomers, fibrils, and DFDNB stabilized oligomers as a function of time. For both B and C, the error bars represent standard deviation across three independent experiments.

### Oligomer structural flexibility plays a greater role than altered charge in oligomer binding to lipid membranes

Treatment with DFDNB alters oligomers in two ways: 1) creating covalent crosslinks that restricts conformational flexibility and 2) removing the positive charge of lysine residues. To evaluate the relative roles of these changes in inhibiting oligomers binding to lipids, truncated Nt17 peptides (no polyQ, Figure 7A) were incorporated into htt-exon1(20Q) oligomers. Nt17 peptides retard htt fibril formation by incorporating into oligomers and creating space between polyQ domains.^34, 57^ While this strategy slows fibril formation, fibrils still eventually form from htt proteins with expanded polyQ domains, which is why htt-exon1 with only 20 repeat glutamine residues were used. To reduce the charge associated with lysine resides, an Nt17 peptides with all three lysine residues mutated to glutamine residues (K6Q, K9Q, K15Q) were also used. These mixed oligomers still retain some lysine associated charge due to the presence of unaltered htt-exon1(20Q). The resulting oligomers can also be treated with DFDNB.

**Figure 7.**
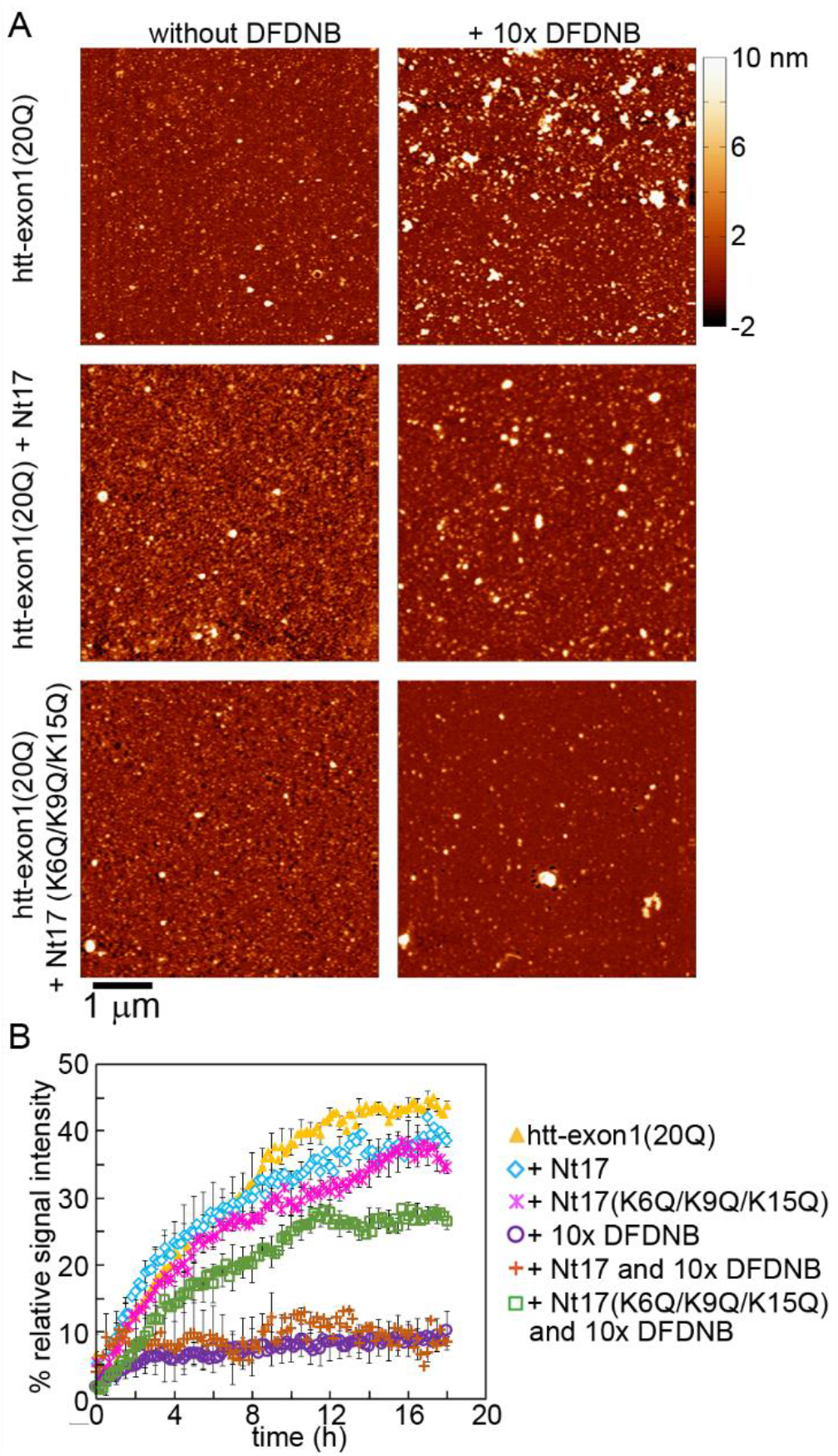
(A) Representative AFM images of htt-exon1(20Q) oligomers that were formed in the absence and presence of Nt17 peptides (both WT and K6Q/K9Q/K15Q). These oligomers were either untreated (without DFDNB) and imaged after 3 h of incubation, or treated with 10x DFDNB and imaged after 24 h of additional incubation. (B) The relative affinity of various preparations of htt-exon1(20Q) oligomers for lipid membranes measured by a PDA/TBLE lipid binding assay.

Htt-exon1(20Q) was incubated at 20 µM alone and with either 20 µM wild type Nt17 or Nt17(K6Q, K9Q, K15Q). After 3 h, each incubation was divided, with half treated with 10x DFDNB and the other half diluted with a corresponding dose of buffer. Untreated and crosslinked htt-exon1(20Q) oligomers were incubated an additional 24 h and evaluated by AFM (Figures 7B). Large, stable population of oligomers (without fibrils) similar in size were present under all conditions. Oligomer membrane activity was determined by a PDA/TBLE lipid binding assay (10 µM of htt-exon1(20Q), Figure 7C). Similar to experiments with htt-exon1(46Q), oligomers comprised of only htt-exon1(20Q) invoked a steady relative CR (maximum of ∼46%) that was completely inhibited by DFDNB treatment. Incorporating Nt17 peptides (both WT and K6Q/KQ9/K15Q) into htt-exon1(20Q) slightly reduced the CR (∼40%), potentially due to the missing polyQ domains. DFDNB treatment of htt-exon1(46Q) oligomers containing WT Nt17 strongly inhibits their ability to bind vesicles in a manner similar to that invoked by DFDNB treatment of pure htt-exon1(20Q) oligomers, indicating that if lysine residues are available that DFDNB will strongly inhibit oligomer lipid-binding. In contrast, DFDNB treatment of htt-exon1(20Q) oligomers with Nt17(K6Q/K9Q/K15Q) only slightly reduced the ability of oligomers to bind vesicles (max CR 28%). These mutations reduced the number of potential crosslinking sites within an oligomer, reducing the induced restrictions on structural flexibility. However, the charge of the available lysine residues is still altered by DFDNB. Collectively, these observations suggest that restrictions in oligomer structural flexibility play a greater role in inhibiting the ability of htt oligomers to bind lipid membrane compared with removing the charge associated with lysine residues.

### Crosslinking reduces oligomer toxicity

To determine if crosslinking altered htt oligomer toxicity, an MTT cytotoxicity assay was performed on N2a neuronal cultured cells (Figure 8). Cells were exposed to freshly prepared htt-exon1(46Q) oligomers, DFDNB stabilized oligomers (5× and 20× treatments), and a DFDNB vehicle control. The unmodified and crosslinked oligomers were added to the culture medium in doses ranging from 0.2-5 μM final concentration. DFDNB concentration in vehicle controls ranged from ∼0.012-0.3 µM based on the amount needed to crosslink oligomers at 20× and the final volume added to each well. Complicating the assay, DFDNB vehicle exhibited comparable toxicity to htt oligomers. Exposure to 0.03 µM or higher DFDNB vehicle (corresponding to the 0.5 µM or higher htt treatment) reduced cell viability to less than 23 % relative to control (N2a cells not exposed to htt or DFDNB); however, cell viability was 67.3 ± 4.0 % with exposure to 0.012 μM DFDNB (corresponding to the 0.2 µM htt treatment).

**Figure 8.**
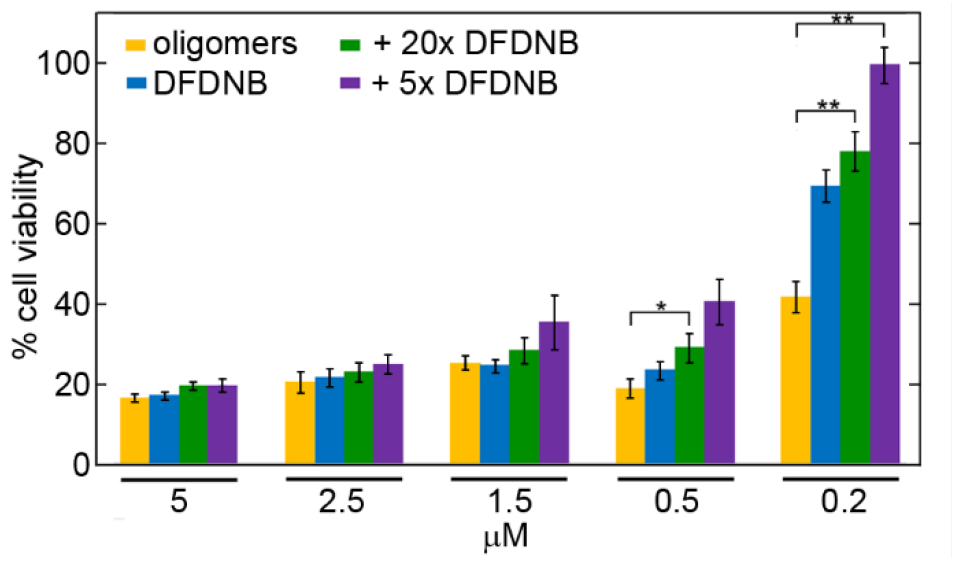
MTT toxicity assay for the N2A cells exposed to htt-exon1(46Q) oligomers and crosslinked oligomers with 5× and 20× DFDNB treatment. * indicates p < 0.05, ** indicates p < 0.01 based on a t test (n = 3). Error bars represent standard error of the mean (SEM).

Unmodified htt-exon1(46Q) oligomers were toxic to N2A cells in a dose-dependent manner. Cell viability was 15.8 ± 1.0 % relative to control with 5 µM htt oligomers, and this increased to 40.4 ± 3.6 % upon exposure to unmodified oligomers at 0.2 μM. With stabilized oligomers treated with 5× or 20× DFDNB, there was not a significant change in cell viability at doses greater than 1.5 µM; although, there is a trend of increased viability relative to control with decreasing concentration. The issue in interpreting this result is that toxicity derives from both untreated htt oligomers and residual DFDNB. With 0.5 µM stabilized oligomers, there is a statistically significant (p < 0.05) increase in cell viability (39.1 ± 5.5% relative to control) with 5× DFDNB stabilized oligomers, and reduced toxicity associated with 20× DFDNB treated oligomers that is not quite statistically significant. Both 5× (96.8 ± 4.3% cell viability) and 20× (75.7 ± 4.8% cell viability) treated DFDNB oligomers are significantly less toxic (p < 0.01 for both compared to unmodified oligomers) at 0.2 µM htt. In short, unmodified htt (no DFDNB added) considerably reduced cell viability with doses as low as 0.2 µM. Vehicle control containing DFDNB (but no htt) reduced cell viability as well, potentially masking the impact of DFDNB treatment on htt oligomer toxicity at high protein concentrations. However, as the DFDNB was effectively diluted out with smaller doses of htt, DFDNB treated oligomers became less toxic relative to htt control. This suggests that crosslinking reduces oligomer toxicity. In addition, crosslinking removes free DFDNB from solution, effectively reducing DFDNB toxicity. This suggests a link between oligomer membrane activity and toxicity.

## DISCUSSION

While damage to various subcellular membranes is evident in HD,^5, 22-28^ little is known regarding how specific aggregate species interact with lipid membranes. Here, the membrane activity of various aggregation states of htt was determined, implicating oligomers as potent membrane disruptors. Fibrillization dramatically reduced the ability of htt to directly interact with membranes. Due to the role of lysine residues in aggregation and oligomer structure,^47^ a known lysine specific crosslinking agent, DFDNB, was employed to further investigate the oligomer/membrane interaction. Rather than stabilizing oligomers, small doses of DFDNB accelerated fibrillization, but with larger doses oligomer stabilization was achieved. However, stabilization precluded membrane activity and reduced associated cytotoxicity of oligomers. Incorporation of stabilizing Nt17 peptides into oligomers with and without mutated lysine residues pointed to a larger role of structural flexibility for htt oligomers to bind membranes compared to loss of charge associated with treatment with DFDNB. Collectively, this suggests that conformational flexibility underlies the membrane activity of htt oligomers and that targeting structural features of htt oligomers represents a potential strategy to block related toxicity.

Despite years of studies, the precise molecular mechanism of htt aggregation and its relation to cellular toxicity is not completely understood. The temporal emergence of various aggregate states has been linked to numerous pathological events associated with mutant htt. The complexity of htt aggregation, along with the metastability of many aggregate intermediates, has made precise assignment and mechanistic understanding of toxic gain of function to specific aggregate species difficult. Due to these issues being common across many amyloid diseases, a variety of crosslinking methods have been employed to stabilize oligomers for structural and functional analyses.^55, 58, 59^ Our results have several implications for these strategies. First, crosslinking can alter the biophysical properties of oligomers, altering their exogenous interactions. Here, it was demonstrated that altering conformational flexibility impacts the membrane activity of htt oligomers. Second, this impact on oligomer membrane activity suggests that crosslinking or otherwise interfering with conformational flexibility of oligomers may represent a therapeutic strategy. However, such an approach would require a crosslinking method that is more selective than DFDNB or other commonly used crosslinking methods.

Lipid membranes modulate htt-exon1 aggregation.^46, 60^ However, fibril formation can be either stimulated^25, 43, 46, 61^ or reduced^43-45, 61, 62^ by membranes depending on the precise lipid composition. Here, aggregation state further modifies the htt/lipid interaction with oligomers being the most aggressive. This is consistent with htt oligomers inducing mitochondrial fragmentation and ER stress.^27, 63^ However, larger aggregate species have previously been implicated in membrane dysfunction. For example, htt inclusions impinge upon the ER^22^ and the nuclear envelope ^23^ leading to altered membrane dynamics, deformation, and disruption. While these studies associated htt fibrils/inclusions with changes in ER membranes or the nuclear envelope, this may represent a late stage marker of earlier events, as it is plausible that subtle alterations in these membranes are induced by earlier htt aggregates or even the aggregation process. Such a notion is supported by the ability of un-aggregated htt to invoke localized morphological and mechanical changes on lipid membranes^39, 56, 64^ and our observation reported here that fibrils have significantly reduced membrane activity.

A plausible mechanism for the enhanced ability of oligomers to bind membranes involves availability of induced α-helical structure of Nt17 (Figure 9A). Nt17-mediated htt membrane binding involves four basic steps: approach, reorganization, anchoring, and insertion.^65, 66^ In short, intrinsically disordered Nt17 stabilizes interactions with membranes by forming an amphipathic α-helix. The time required for reorganizing into that α-helix provides a window for dissociate from the lipid bilayer.^48^ In solution, Nt17 exists in a concentration-dependent equilibrium between an intrinsically disordered monomeric state and these α-helical oligomers.^35, 67^ However, Nt17 packed into oligomers has already adopted α-helical structure.^37^ Stabilization of the Nt17 α-helix by oligomerization may bypass the reorganization step, enhancing the membrane activity of oligomers. Such a mechanism requires two assumptions: 1) that preformed α-helixes bind membranes faster and 2) that oligomer structure remains flexible enough for Nt17 α-helices to interact with lipids. Previous computational studies support the first assumption as preconfiguring Nt17 into α-helixes accelerates binding between Nt17 and lipids.^68^ The second assumption is supported by crosslinking reducing oligomer membrane activity and MD simulations demonstrating that tetrameric Nt17 oligomers are structurally dynamic.

**Figure 9.**
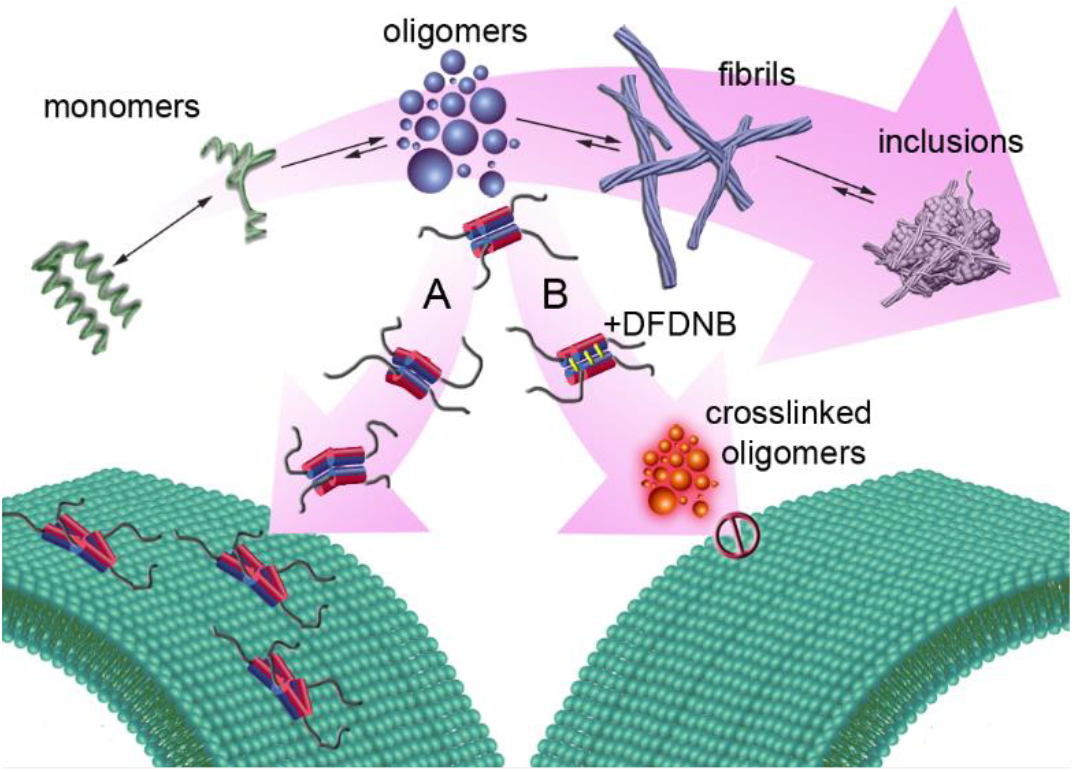
The proposed mechanism for interaction of (A) oligomers interact readily with lipid membranes and (B) lysine-crosslinked oligomers leads to diminished interaction with lipid membrane

Reduced membrane activity of fibrils may be linked to the availability of Nt17. Htt fibrils are comprised of a rigid amyloid core of polyQ domains surrounded by both the Nt17 and proline-rich domains (PRD).^69-71^ While there are variations across studies, the PRD is the more flexible and solvent exposed of the two flanking regions in fibrils.^70, 72-74^ In reported fibril structures, the dynamics and conformation of Nt17 varies, ranging from being completely buried in a packed structure,^73^ to dynamic heterogeneity, ^72^ or to being helical with partial immobilization by dense packing on the fibril surface.^38, 70, 75^ Decreased membrane activity of htt fibrils suggests that Nt17 is either not accessible within mature fibrils or at least is partially bracketed with non-helical structure that restrict initial binding. Additionally, MD studies suggest that polyQ forms hydrogen bonds with phospholipid head groups, providing a scaffold for further insertion of Nt17 into the membrane core.^66^ Thus, dense packing of polyQ in the amyloid core can also reduce membrane binding of fibrils.

While high doses of DFDNB stabilized oligomers, it was surprising that low doses accelerated fibrillization. As both intra and intermolecular crosslinking occurs with DFDNB, such a strategy results in an inherent complexity of crosslinking products.^76, 77^ Low concentration of DFDNB relative to the target substrate promotes intramolecular crosslinking, whereas, efficiency of intermolecular crosslinking is enhanced with higher DFDNB doses.^78^ With three lysine residues in Nt17, lower doses (< 10 µM) of DFDNB result in an excess of target residues, promoting intramolecular crosslinking. Based on acetylation studies, lysine 9 is the most readily available lysine for modification.^48, 79^ Lysine 9 is also not well-positioned for intermolecular crosslinks based on available structures of htt oligomers.^37, 47^ Based on spacing in the α-helix with respect to Lysine 6 and the flexibility associated with lysine 15, intramolecular crosslinks with lysine 9 are plausible with both of these residues. These crosslinks would have a greater impact on helix stability rather than relative motions of helices with respect to each other in oligomers. A small amount of these intramolecular crosslinks could push the equilibrium between disordered Nt17 monomers and higher order multimers toward oligomerization without restricting conformational freedom needed for initiating fibrillization. With larger DFDNB treatments, saturation of available lysine residues would result in a combination of intramolecular and intermolecular crosslinks, stabilizing oligomers and limiting the dynamic behavior required for fibril nucleation. While a clear consensus of the fate of Nt17 in fibrils is lacking,^80^ forming the densely packed polyQ amyloid core likely requires some conformational flexibility.

With low doses of DFDNB that promoted fibril formation, the partial reduction of membrane activity by non-stabilized oligomers may encompass a variety of mechanisms. By promoting fibrils, the highly membrane active oligomer population would be reduced as membrane inactive fibrils form. With higher doses of DFDNB, conformational flexibility of α-helical bundles is compromised by intermolecular DFDNB-mediated crosslinking of lysine residues, and the membrane activity of stabilized oligomers is completely inhibited. (Figure 9B). Altering charge associated with lysine residues had a smaller impact in comparison to hindered structural flexibility. This is consistent with extensive lysine acetylation, which also removes the positive charge, reducing membrane activity of htt-exon1 by ∼40%, ^48^ and a generic role of lysine acetylation modulating membrane affinity of proteins.^81^ Importantly, the loss of membrane activity of oligomers coincides with a reduction in cellular toxicity, suggesting that targeting structural flexibility of oligomers may provide therapeutic benefit.

## REFERENCES

(1) Gatchel, J. R.; Zoghbi, H. Y. Diseases of Unstable Repeat Expansion: Mechanisms and Common Principles. Nature Reviews Genetics 2005, 6 (10), 743–755. DOI: 10.1038/nrg1691.

(2) Adegbuyiro, A.; Sedighi, F.; Pilkington, A. W.; Groover, S.; Legleiter, J. Proteins Containing Expanded Polyglutamine Tracts and Neurodegenerative Disease. Biochemistry 2017, 56 (9), 1199–1217. DOI: 10.1021/acs.biochem.6b00936.

(3) The Huntington’s Disease Collobartive Research Group. A novel gene containing a trinucleotide repeat that is exanded and unstable on Huntingtons-disease chromosomes Cell 1993, 72 (6), 971–983. DOI: 10.1016/0092-8674(93)90585-e.

(4) Davies, S. W.; Turmaine, M.; Cozens, B. A.; DiFiglia, M.; Sharp, A. H.; Ross, C. A.; Scherzinger, E.; Wanker, E. E.; Mangiarini, L.; Bates, G. P. Formation of neuronal intranuclear inclusions underlies the neurological dysfunction in mice transgenic for the HD mutation. Cell 1997, 90 (3), 537–548. DOI: 10.1016/s0092-8674(00)80513-9.

(5) Gutekunst, C. A.; Li, S. H.; Yi, H.; Mulroy, J. S.; Kuemmerle, S.; Jones, R.; Rye, D.; Ferrante, R. J.; Hersch, S. M.; Li, X. J. Nuclear and neuropil aggregates in Huntington’s disease: Relationship to neuropathology. J Neurosci 1999, 19 (7), 2522–2534.

(6) Wagner, A. S.; Politi, A. Z.; Ast, A.; Bravo-Rodriguez, K.; Baum, K.; Buntru, A.; Strempel, N. U.; Brusendorf, L.; Hänig, C.; Boeddrich, A.; et al. Self-assembly of Mutant Huntingtin Exon-1 Fragments into Large Complex Fibrillar Structures Involves Nucleated Branching. J Mol Biol 2018, 430 (12), 1725–1744. DOI: 10.1016/j.jmb.2018.03.017.

(7) Sahoo, B.; Arduini, I.; Drombosky, K. W.; Kodali, R.; Sanders, L. H.; Greenamyre, J. T.; Wetzel, R. Folding Landscape of Mutant Huntingtin Exon1: Diffusible Multimers, Oligomers and Fibrils, and No Detectable Monomer. PLoS One 2016, 11 (6). DOI: 10.1371/journal.pone.0155747.

(8) Legleiter, J.; Mitchell, E.; Lotz, G. P.; Sapp, E.; Ng, C.; DiFiglia, M.; Thompson, L. M.; Muchowski, P. J. Mutant Huntingtin Fragments Form Oligomers in a Polyglutamine Length-dependent Manner in Vitro and in Vivo. J Biol Chem 2010, 285 (19), 14777–14790. DOI: 10.1074/jbc.M109.093708.

(9) Landrum, E.; Wetzel, R. Biophysical Underpinnings of the Repeat Length Dependence of Polyglutamine Amyloid Formation. J Biol Chem 2014, 289 (15), 10254–10260. DOI: 10.1074/jbc.C114.552943.

(10) Sahl, S. J.; Lau, L.; Vonk, W. I. M.; Weiss, L. E.; Frydman, J.; Moerner, W. E. Delayed emergence of subdiffraction-sized mutant huntingtin fibrils following inclusion body formation. Q Rev Biophys 2015, 49. DOI: 10.1017/s0033583515000219.

(11) Sahl, S. J.; Weiss, L. E.; Duim, W. C.; Frydman, J.; Moerner, W. E. Cellular Inclusion Bodies of Mutant Huntingtin Exon 1 Obscure Small Fibrillar Aggregate Species. Sci Rep 2012, 2. DOI: 10.1038/srep00895.

(12) Saudou, F.; Finkbeiner, S.; Devys, D.; Greenberg, M. E. Huntingtin acts in the nucleus to induce apoptosis but death does not correlate with the formation of intranuclear inclusions. Cell 1998, 95 (1), 55–66. DOI: 10.1016/s0092-8674(00)81782-1.

(13) Arrasate, M.; Mitra, S.; Schweitzer, E. S.; Segal, M. R.; Finkbeiner, S. Inclusion body formation reduces levels of mutant huntingtin and the risk of neuronal death. Nature 2004, 431 (7010), 805–810. DOI: 10.1038/nature02998.

(14) Olshina, M. A.; Angley, L. M.; Ramdzan, Y. M.; Tang, J.; Bailey, M. F.; Hill, A. F.; Hatters, D. M. Tracking Mutant Huntingtin Aggregation Kinetics in Cells Reveals Three Major Populations That Include an Invariant Oligomer Pool. J Biol Chem 2010, 285 (28), 21807–21816. DOI: 10.1074/jbc.M109.084434.

(15) Lajoie, P.; Snapp, E. L. Formation and Toxicity of Soluble Polyglutamine Oligomers in Living Cells. PLoS One 2010, 5 (12). DOI: 10.1371/journal.pone.0015245.

(16) Lu, M.; Banetta, L.; Young, L. J.; Smith, E. J.; Bates, G. P.; Zaccone, A.; Schierle, G. S. K.; Tunnacliffe, A.; Kaminski, C. F. Live-cell super-resolution microscopy reveals a primary role for diffusion in polyglutamine-driven aggresome assembly. J Biol Chem 2019, 294 (1), 257–268. DOI: 10.1074/jbc.RA118.003500.

(17) Nagai, Y.; Inui, T.; Popiel, H. A.; Fujikake, N.; Hasegawa, K.; Urade, Y.; Goto, Y.; Naiki, H.; Toda, T. A toxic monomeric conformer of the polyglutamine protein. Nat Struct Mol Biol 2007, 14 (4), 332–340. DOI: 10.1038/nsmb1215.

(18) Nucifora, L. G.; Burke, K. A.; Feng, X.; Arbez, N.; Zhu, S.; Miller, J.; Yang, G.; Ratovitski, T.; Delannoy, M.; Muchowski, P. J.; et al. Identification of Novel Potentially Toxic Oligomers Formed in Vitro from Mammalian-derived Expanded huntingtin Exon-1 Protein. J Biol Chem 2012, 287 (19), 16017–16028. DOI: 10.1074/jbc.M111.252577.

(19) Kim, Y. E.; Hosp, F.; Frottin, F.; Ge, H.; Mann, M.; Hayer-Hartl, M.; Hartl, F. U. Soluble Oligomers of PolyQ-Expanded Huntingtin Target a Multiplicity of Key Cellular Factors. Mol Cell 2016, 63 (6), 951–964. DOI: 10.1016/j.molcel.2016.07.022.

(20) Drombosky, K. W.; Rode, S.; Kodali, R.; Jacob, T. C.; Palladino, M. J.; Wetzel, R. Mutational analysis implicates the amyloid fibril as the toxic entity in Huntington’s disease. Neurobiol Dis 2018, 120, 126–138. DOI: 10.1016/j.nbd.2018.08.019.

(21) Pieri, L.; Madiona, K.; Bousset, L.; Melki, R. Fibrillar alpha-Synuclein and Huntingtin Exon 1 Assemblies Are Toxic to the Cells. Biophys J 2012, 102 (12), 2894–2905. DOI: 10.1016/j.bpj.2012.04.050.

(22) Bäuerlein, F. J. B.; Saha, I.; Mishra, A.; Kalemanov, M.; Martinez-Sanchez, A.; Klein, R.; Dudanova, I.; Hipp, M. S.; Hartl, F. U.; Baumeister, W.; et al. In Situ Architecture and Cellular Interactions of PolyQ Inclusions. Cell 2017, 171 (1), 179-+. DOI: 10.1016/j.cell.2017.08.009.

(23) Liu, K.-Y.; Shyu, Y.-C.; Barbaro, B. A.; Lin, Y.-T.; Chern, Y.; Thompson, L. M.; Shen, C.-K. J.; Marsh, J. L. Disruption of the nuclear membrane by perinuclear inclusions of mutant huntingtin causes cell-cycle re-entry and striatal cell death in mouse and cell models of Huntington’s disease. Hum Mol Genet 2015, 24 (6), 1602–1616. DOI: 10.1093/hmg/ddu574.

(24) Harjes, P.; Wanker, E. E. The hunt for huntingtin function: interaction partners tell many different stories. Trends Biochem Sci 2003, 28 (8), 425–433. DOI: 10.1016/s0968-0004(03)00168-3.

(25) Orr, H. T.; Chung, M. Y.; Banfi, S.; Kwiatkowski, T. J.; Servadio, A.; Beaudet, A. L.; McCall, A. E.; Duvick, L. A.; Ranum, L. P. W.; Zoghbi, H. Y. EXPANSION OF AN UNSTABLE TRINUCLEOTIDE CAG REPEAT IN SPINOCEREBELLAR ATAXIA TYPE-1. Nat Genet 1993, 4 (3), 221–226. DOI: 10.1038/ng0793-221.

(26) Panov, A. V.; Gutekunst, C.-A.; Leavitt, B. R.; Hayden, M. R.; Burke, J. R.; Strittmatter, W. J.; Greenamyre, J. T. Early mitochondrial calcium defects in Huntington’s disease are a direct effect of polyglutamines. Nat Neurosci 2002, 5 (8), 731–736. DOI: 10.1038/nn884.

(27) Shirendeb, U.; Reddy, A. P.; Manczak, M.; Calkins, M. J.; Mao, P.; Tagle, D. A.; Reddy, P. H. Abnormal mitochondrial dynamics, mitochondrial loss and mutant huntingtin oligomers in Huntington’s disease: implications for selective neuronal damage. Hum Mol Genet 2011, 20 (7), 1438–1455. DOI: 10.1093/hmg/ddr024.

(28) Ueda, M.; Li, S.; Itoh, M.; Wang, M.-x.; Hayakawa, M.; Islam, S.; Tana Nakagawa, K.; Chen, H.; Nakagawa, T. Expanded polyglutamine embedded in the endoplasmic reticulum causes membrane distortion and coincides with Bax insertion. Biochem Bioph Res Co 2016, 474 (2), 259–263. DOI: 10.1016/j.bbrc.2016.04.034.

(29) Ho, C. S.; Khadka, N. K.; She, F.; Cai, J.; Pan, J. Polyglutamine aggregates impair lipid membrane integrity and enhance lipid membrane rigidity. BBA - Biomembranes 2016, 1858 (4), 661–670. DOI: 10.1016/j.bbamem.2016.01.016.

(30) Qin, Z. H.; Wang, Y. M.; Sapp, E.; Cuiffo, B.; Wanker, E.; Hayden, M. R.; Kegel, K. B.; Aronin, N.; DiFiglia, M. Huntingtin bodies sequester vesicle-associated proteins by a polyproline-dependent interaction. J Neurosci 2004, 24 (1), 269–281. DOI: 10.1523/jneurosci.1409-03.2004.

(31) Gasset-Rosa, F.; Chillon-Marinas, C.; Goginashvili, A.; Atwal, R. S.; Artates, J. W.; Tabet, R.; Wheeler, V. C.; Bang, A. G.; Cleveland, D. W.; Lagier-Tourenne, C. Polyglutamine-Expanded Huntingtin Exacerbates Age-Related Disruption of Nuclear Integrity and Nucleocytoplasmic Transport. Neuron 2017, 94 (1), 48-57.e44. DOI: 10.1016/j.neuron.2017.03.027.

(32) Riguet, N.; Mahul-Mellier, A.-L.; Maharjan, N.; Burtscher, J.; Patin, A.; Croisier, M.; Knott, G.; Reiterer, V.; Farhan, H.; Lashuel, H. A. Disentangling the sequence, cellular and ultrastructural determinants of Huntingtin nuclear and cytoplasmic inclusion formation. Cold Spring Harbor Laboratory: 2020.

(33) Jayaraman, M.; Kodali, R.; Sahoo, B.; Thakur, A. K.; Mayasundari, A.; Mishra, R.; Peterson, C. B.; Wetzel, R. Slow Amyloid Nucleation via alpha-Helix-Rich Oligomeric Intermediates in Short Polyglutamine-Containing Huntingtin Fragments. J Mol Biol 2012, 415 (5), 881–899. DOI: 10.1016/j.jmb.2011.12.010.

(34) Mishra, R.; Jayaraman, M.; Roland, B. P.; Landrum, E.; Fullam, T.; Kodali, R.; Thakur, A. K.; Arduini, I.; Wetzel, R. Inhibiting the Nucleation of Amyloid Structure in a Huntingtin Fragment by Targeting alpha-Helix-Rich Oligomeric Intermediates. J Mol Biol 2012, 415 (5), 900–917. DOI: 10.1016/j.jmb.2011.12.011.

(35) Thakur, A. K.; Jayaraman, M.; Mishra, R.; Thakur, M.; Chellgren, V. M.; Byeon, I.-J. L.; Anjum, D. H.; Kodali, R.; Creamer, T. P.; Conway, J. F.; et al. Polyglutamine disruption of the huntingtin exon 1 N terminus triggers a complex aggregation mechanism. Nat Struct Mol Biol 2009, 16 (4), 380–389. DOI: 10.1038/nsmb.1570.

(36) Michalek, M.; Salnikov, E. S.; Werten, S.; Bechinger, B. Membrane Interactions of the Amphipathic Amino Terminus of Huntingtin. Biochemistry 2013, 52 (5), 847–858. DOI: 10.1021/bi301325q.

(37) Kotler, S. A.; Tugarinov, V.; Schmidt, T.; Ceccon, A.; Libich, D. S.; Ghirlando, R.; Schwieters, C. D.; Clore, G. M. Probing initial transient oligomerization events facilitating Huntingtin fibril nucleation at atomic resolution by relaxation-based NMR. P Nat Acad Sci USA 2019, 116 (9), 3562–3571. DOI: 10.1073/pnas.1821216116.

(38) Mishra, R.; Hoop, C. L.; Kodali, R.; Sahoo, B.; van der Wel, P. C. A.; Wetzel, R. Serine Phosphorylation Suppresses Huntingtin Amyloid Accumulation by Altering Protein Aggregation Properties. J Mol Biol 2012, 424 (1-2), 1–14. DOI: 10.1016/j.jmb.2012.09.011.

(39) Burke, K. A.; Kauffman, K. J.; Umbaugh, C. S.; Frey, S. L.; Legleiter, J. The Interaction of Polyglutamine Peptides with Lipid Membranes Is Regulated by Flanking Sequences Associated with Huntingtin. J Biol Chem 2013, 288 (21), 14993–15005. DOI: 10.1074/jbc.M112.446237.

(40) Michalek, M.; Salnikov, E. S.; Bechinger, B. Structure and Topology of the Huntingtin 1-17 Membrane Anchor by a Combined Solution and Solid-State NMR Approach. Biophys J 2013, 105 (3), 699–710. DOI: 10.1016/j.bpj.2013.06.030.

(41) Chaibva, M.; Burke, K. A.; Legleiter, J. Curvature Enhances Binding and Aggregation of Huntingtin at Lipid Membranes. Biochemistry 2014, 53 (14), 2355–2365. DOI: 10.1021/bi401619q.

(42) Atwal, R. S.; Xia, J.; Pinchev, D.; Taylor, J.; Epand, R. M.; Truant, R. Huntingtin has a membrane association signal that can modulate huntingtin aggregation, nuclear entry and toxicity. Hum Mol Genet 2007, 16 (21), 2600–2615. DOI: 10.1093/hmg/ddm217.

(43) Beasley, M.; Groover, S.; Valentine, S. J.; Legleiter, J. Lipid headgroups alter huntingtin aggregation on membranes. BBA-Biomembranes 2021, 1863 (1), 183497. DOI: https://doi.org/10.1016/j.bbamem.2020.183497.

(44) Chaibva, M.; Gao, X.; Jain, P.; Campbell, W. A.; Frey, S. L.; Legleiter, J. Sphingomyelin and GM1 Influence Huntingtin Binding to, Disruption of, and Aggregation on Lipid Membranes. ACS Omega 2018, 3 (1), 273–285. DOI: 10.1021/acsomega.7b01472.

(45) Gao, X.; Campbell, W. A.; Chaibva, M.; Jain, P.; Leslie, A. E.; Frey, S. L.; Legleiter, J. Cholesterol Modifies Huntingtin Binding to, Disruption of, and Aggregation on Lipid Membranes. Biochemistry 2016, 55 (1), 92–102. DOI: 10.1021/acs.biochem.5b00900.

(46) Pandey, N. K.; Isas, J. M.; Rawat, A.; Lee, R. V.; Langen, J.; Pandey, P.; Langen, R. The 17-residue-long N terminus in huntingtin controls stepwise aggregation in solution and on membranes via different mechanisms. J Biol Chem 2018, 293 (7), 2597–2605. DOI: 10.1074/jbc.M117.813667.

(47) Arndt, J. R.; Brown, R. J.; Burke, K. A.; Legleiter, J.; Valentine, S. J. Lysine residues in the N-terminal huntingtin amphipathic alpha-helix play a key role in peptide aggregation. J Mass Spectrom 2015, 50 (1), 117–126. DOI: 10.1002/jms.3504.

(48) Chaibva, M.; Jawahery, S.; Pilkington, A. W.; Arndt, J. R.; Sarver, O.; Valentine, S.; Matysiak, S.; Legleiter, J. Acetylation within the First 17 Residues of Huntingtin Exon 1 Alters Aggregation and Lipid Binding. Biophys J 2016, 111 (2), 349–362. DOI: 10.1016/j.bpj.2016.06.018.

(49) Sedighi, F.; Adegbuyiro, A.; Legleiter, J. SUMOylation Prevents Huntingtin Fibrillization and Localization onto Lipid Membranes. ACS Chem Neurosci 2020, 11 (3), 328–343. DOI: 10.1021/acschemneuro.9b00509.

(50) Muchowski, P. J.; Schaffar, G.; Sittler, A.; Wanker, E. E.; Hayer-Hartl, M. K.; Hartl, F. U. Hsp70 and Hsp40 chaperones can inhibit self-assembly of polyglutamine proteins into amyloid-like fibrils. P Natl Acad Sci USA 2000, 97 (14), 7841–7846. DOI: 10.1073/pnas.140202897.

(51) Burke, K. A.; Godbey, J.; Legleiter, J. Assessing mutant huntingtin fragment and polyglutamine aggregation by atomic force microscopy. Methods 2011, 53 (3), 275–284. DOI: 10.1016/j.ymeth.2010.12.028.

(52) Beasley, M.; Stonebraker, A. R.; Legleiter, J. Normalizing polydiacetylene colorimetric assays of vesicle binding across lipid systems. Anal Biochem 2020, 609. DOI: 10.1016/j.ab.2020.113864.

(53) Kolusheva, S.; Boyer, L.; Jelinek, R. A colorimetric assay for rapid screening of antimicrobial peptides. Nat Biotech 2000, 18 (2), 225–227. DOI: 10.1038/72697.

(54) Zheng, F.; Wu, Z.; Chen, Y. A quantitative method for the measurement of membrane affinity by polydiacetylene-based colorimetric assay. Anal Biochem 2012, 420 (2), 171–176. DOI: 10.1016/j.ab.2011.09.026.

(55) Cline, E. N.; Das, A.; Bicca, M. A.; Mohammad, S. N.; Schachner, L. F.; Kamel, J. M.; Dinunno, N.; Weng, A.; Paschall, J. D.; Bu, R. L.; et al. A novel crosslinking protocol stabilizes amyloid β oligomers capable of inducing Alzheimer’s‐associated pathologies. J Neurochem 2019, 148 (6), 822–836. DOI: 10.1111/jnc.14647.

(56) Burke, K. A.; Hensal, K. M.; Umbaugh, C. S.; Chaibva, M.; Legleiter, J. Huntingtin disrupts lipid bilayers in a polyQ-length dependent manner. BBA - Biomembranes 2013, 1828 (8), 1953–1961. DOI: 10.1016/j.bbamem.2013.04.025.

(57) Arndt, J. R.; Chaibva, M.; Beasley, M.; Karanji, A. K.; Kondalaji, S. G.; Khakinejad, M.; Sarver, O.; Legleiter, J.; Valentine, S. J. Nucleation Inhibition of Huntingtin Protein (htt) by Polyproline PPII Helices: A Potential Interaction with the N-Terminal alpha-Helical Region of Htt. Biochemistry 2020, 59 (4), 436–449. DOI: 10.1021/acs.biochem.9b00689.

(58) Bitan, G.; Lomakin, A.; Teplow, D. B. Amyloid β-Protein Oligomerization. J Biol Chem 2001, 276 (37), 35176–35184. DOI: 10.1074/jbc.m102223200.

(59) Williams, T. L.; Serpell, L. C.; Urbanc, B. Stabilization of native amyloid β-protein oligomers by Copper and Hydrogen peroxide Induced Cross-linking of Unmodified Proteins (CHICUP). BBA - Proteins and Proteomics 2016, 1864 (3), 249–259. DOI: 10.1016/j.bbapap.2015.12.001.

(60) Levy, G. R.; Shen, K.; Gavrilov, Y.; Smith, P. E. S.; Levy, Y.; Chan, R.; Frydman, J.; Frydman, L. Huntingtin’s N-Terminus Rearrangements in the Presence of Membranes: A Joint Spectroscopic and Computational Perspective. ACS Chem Neurosci 2019, 10 (1), 472–481. DOI: 10.1021/acschemneuro.8b00353.

(61) Beasley, M.; Frazee, N.; Groover, S.; Valentine, S. J.; Mertz, B.; Legleiter, J. Physicochemical Properties Altered by the Tail Group of Lipid Membranes Influence Huntingtin Aggregation and Lipid Binding. Phys Chem B 2022, 126 (16), 3067–3081. DOI: 10.1021/acs.jpcb.1c10254.

(62) Adegbuyiro, A.; Sedighi, F.; Jain, P.; Pinti, M. V.; Siriwardhana, C.; Hollander, J. M.; Legleiter, J. Mitochondrial membranes modify mutant huntingtin aggregation. BBA-Biomembranes 2021, 1863 (10), 183663. DOI: https://doi.org/10.1016/j.bbamem.2021.183663.

(63) Leitman, J.; Hartl, F. U.; Lederkremer, G. Z. Soluble forms of polyQ-expanded huntingtin rather than large aggregates cause endoplasmic reticulum stress. Nat Commun 2013, 4. DOI: 10.1038/ncomms3753.

(64) Burke, K. A.; Yates, E. A.; Legeiter, J. Amyloid-Forming Proteins Alter the Local Mechanical Properties of Lipid Membranes. Biochemistry 2013, 52 (5), 808–817. DOI: 10.1021/bi301070v.

(65) Cote, S.; Binette, V.; Salnikov, E. S.; Bechinger, B.; Mousseau, N. Probing the Huntingtin 1-17 Membrane Anchor on a Phospholipid Bilayer by Using All-Atom Simulations. Biophys J 2015, 108 (5), 1187–1198. DOI: 10.1016/j.bpj.2015.02.001.

(66) Cote, S.; Wei, G.; Mousseau, N. Atomistic mechanisms of huntingtin N-terminal fragment insertion on a phospholipid bilayer revealed by molecular dynamics simulations. Proteins 2014, 82 (7), 1409–1427. DOI: 10.1002/prot.24509.

(67) Kelley, N. W.; Huang, X.; Tam, S.; Spiess, C.; Frydman, J.; Pande, V. S. The Predicted Structure of the Headpiece of the Huntingtin Protein and Its Implications on Huntingtin Aggregation. J Mol Biol 2009, 388 (5), 919–927. DOI: 10.1016/j.jmb.2009.01.032.

(68) Karanji, A. K.; Beasley, M.; Sharif, D.; Ranjbaran, A.; Legleiter, J.; Valentine, S. J. Investigating the interactions of the first 17 amino acid residues of Huntingtin with lipid vesicles using mass spectrometry and molecular dynamics. J Mass Spectrom 2020, 55 (1). DOI: 10.1002/jms.4470.

(69) Hoop, C. L.; Lin, H.-K.; Kar, K.; Magyarfalvi, G.; Lamley, J. M.; Boatz, J. C.; Mandal, A.; Lewandowski, J. R.; Wetzel, R.; van der Wel, P. C. A. Huntingtin exon 1 fibrils feature an interdigitated beta-hairpin-based polyglutamine core. P Natl Acad Sci USA 2016, 113 (6), 1546–1551. DOI: 10.1073/pnas.1521933113.

(70) Lin, H.-K.; Boatz, J. C.; Krabbendam, I. E.; Kodali, R.; Hou, Z.; Wetzel, R.; Dolga, A. M.; Poirier, M. A.; van der Wel, P. C. A. Fibril polymorphism affects immobilized non-amyloid flanking domains of huntingtin exon1 rather than its polyglutamine core. Nat Commun 2017, 8. DOI: 10.1038/ncomms15462.

(71) Smith, A. N.; Märker, K.; Piretra, T.; Boatz, J. C.; Matlahov, I.; Kodali, R.; Hediger, S.; Van Der Wel, P. C. A.; De Paëpe, G. Structural Fingerprinting of Protein Aggregates by Dynamic Nuclear Polarization-Enhanced Solid-State NMR at Natural Isotopic Abundance. J Am Chem Soc 2018, 140 (44), 14576–14580. DOI: 10.1021/jacs.8b09002.

(72) Isas, J. M.; Langen, R.; Siemer, A. B. Solid-State Nuclear Magnetic Resonance on the Static and Dynamic Domains of Huntingtin Exon-1 Fibrils. Biochemistry 2015, 54 (25), 3942–3949. DOI: 10.1021/acs.biochem.5b00281.

(73) Bugg, C. W.; Isas, J. M.; Fischer, T.; Patterson, P. H.; Langen, R. Structural Features and Domain Organization of Huntingtin Fibrils. J Biol Chem 2012, 287 (38), 31739–31746. DOI: 10.1074/jbc.M112.353839.

(74) Caulkins, B. G.; Cervantes, S. A.; Isas, J. M.; Siemer, A. B. Dynamics of the Proline-Rich C-Terminus of Huntingtin Exon-1 Fibrils. Phys Chem B 2018, 122 (41), 9507–9515. DOI: 10.1021/acs.jpcb.8b09213.

(75) Sivanandam, V. N.; Jayaraman, M.; Hoop, C. L.; Kodali, R.; Wetzel, R.; van der Wel, P. C. A. The Aggregation-Enhancing Huntingtin N-Terminus Is Helical in Amyloid Fibrils. J Am Chem Soc 2011, 133 (12), 4558–4566. DOI: 10.1021/ja110715f.

(76) Bányai, L.; Patthy, L. Importance of intramolecular interactions in the control of the fibrin affinity and activation of human plasminogen. J Biol Chem 1984, 259 (10), 6466–6471. DOI: 10.1016/s0021-9258(20)82165-6.

(77) Ermácora, M. R.; Nowicki, C.; Wolfenstein-Todel, C.; Santomé, J. A. Identification of intramolecular crosslinks in bovine growth hormone after two-step modification with 1,5-difluoro-2,4-dinitrobenzene. Int J Pept Prot Res 2009, 30 (3), 423–430. DOI: 10.1111/j.1399-3011.1987.tb03350.x.

(78) Nowicki, C.; Wolfenstein-Todel, C.; Santome, J. A. Evidence for the steric proximity of Tyr 174 and Lys 111 in bovine growth hormone. Int J Pept Prot Res 2009, 26 (6), 568–574. DOI: 10.1111/j.1399-3011.1985.tb03213.x.

(79) Cong, X.; Held, J. M.; DeGiacomo, F.; Bonner, A.; Chen, J. M.; Schilling, B.; Czerwieniec, G. A.; Gibson, B. W.; Ellerby, L. M. Mass Spectrometric Identification of Novel Lysine Acetylation Sites in Huntingtin. Mol Cell Proteomics 2011, 10 (10). DOI: 10.1074/mcp.M111.009829.

(80) Matlahov, I.; Van Der Wel, P. C. Conformational studies of pathogenic expanded polyglutamine protein deposits from Huntington’s disease. Exp Biol Med 2019, 244 (17), 1584–1595. DOI: 10.1177/1535370219856620.

(81) Okada, A. K.; Teranishi, K.; Ambroso, M. R.; Isas, J. M.; Vazquez-Sarandeses, E.; Lee, J.-Y.; Melo, A. A.; Pandey, P.; Merken, D.; Berndt, L.; et al. Lysine acetylation regulates the interaction between proteins and membranes. Nat Commun 2021, 12 (1). DOI: 10.1038/s41467-021-26657-2.

